# Functionally redundant control of cardiac hypertrophic signaling by inositol 1,4,5-trisphosphate receptors

**DOI:** 10.1101/075044

**Authors:** M. Iveth Garcia, Anja Karlstaedt, Javier Amione-Guerra, Keith A. Youker, Heinrich Taegtmeyer, Darren Boehning

**Affiliations:** Cell Biology Graduate Program, University of Texas Medical Branch, Galveston, TX 77555; Department of Biochemistry and Molecular Biology, McGovern Medical School at UTHealth, Houston, TX 77030; Department of Internal Medicine, Division of Cardiology, McGovern Medical School at UTHealth, Houston, TX 77030.; Houston Methodist Hospital, Houston, TX 77030.

**Keywords:** IP_3_ Receptor, Calcium, Endothelin-1, Cardiomyocyte, Hypertrophy

## Abstract

Calcium plays an integral role to many cellular processes including contraction, energy metabolism, gene expression, and cell death. The inositol 1,4,5-trisphosphate receptor (IP_3_R) is a calcium channel expressed in cardiac tissue. There are three IP_3_R isoforms encoded by separate genes. In the heart, the IP_3_R-2 isoform is reported to being most predominant with regards to expression levels and functional significance. The functional roles of IP_3_R-1 and IP_3_R-3 in the heart are essentially unexplored despite measureable expression levels. Here we show that all three IP_3_Rs isoforms are expressed in both neonatal and adult rat ventricular cardiomyocytes and in human heart tissue. All three IP_3_R proteins were expressed throughout the cardiomyocyte sarcoplasmic reticulum. Using isoform specific siRNA, we found that expression of all three IP_3_R isoforms are required for hypertrophic signaling downstream of endothelin-1 stimulation. Mechanistically, IP_3_Rs specifically contribute to activation of the hypertrophic program by mediating the positive inotropic effects of endothelin-1 leading to downstream activation of nuclear factor of activated T-cells. Our findings highlight previously unidentified functions for IP_3_R isoforms in the heart with significant implications for hypertrophic signaling in animal models and human disease.

**Significance:** Hypertrophy is an adaptive response to cardiac stress which can lead to arrhythmias and cardiac failure. The peptide hormone endothelin-1(ET-1) is a potent activator of the hypertrophic program in cardiomyocytes. IP_3_R calcium channels are activated downstream of ET-1 during hypertrophy. We now show that all three IP_3_R proteins are essential for hypertrophic signaling downstream of ET-1. Activation of IP_3_Rs did not lead to nuclear-specific calcium transients but instead led to altered contractility ultimately, leading to NFAT activation and activation of the hypertrophic program. These effects were independent of alterations in IP_3_R protein expression levels both in vitro and in the human failing heart. Our results identify a new paradigm in IP_3_R signaling in the heart with relevance to human disease.

## Introduction

Calcium is an essential modulator of a wide variety of cellular functions including cardiomyocyte excitation-contraction coupling (ECC) and gene expression. Cardiomyocyte function is modulated by neuro-hormonal agonists to accommodate cardiac demand. One example is endothelin-1 (ET-1), which is a potent vasoconstrictor that plays an important role in modulating muscle contractility, vascular tone, cardiomyocyte growth, and survival (1, 2). Plasma levels of ET-1 are also increased during pathological conditions such as chronic heart failure, myocardial infarction, cardiac hypertrophy and in hypertension (3, 4). As such, ET-1 has been linked to pathological remodeling of the heart (1, 5). ET-1 signaling is initiated by ET-1 binding to G-protein coupled receptors at the plasma membrane leading to the activation of phospholipase C (PLC). PLC catalyzes the hydrolysis of phosphatidylinositol 4,5-bisphosphate (PIP2) which leads to increased production of the second messengers inositol 1,4,5-trisphosphate (IP_3_) and diacylglycerol. IP_3_ then acts as a second messenger that binds inositol 1,4,5-trisphosphate receptors (IP_3_Rs), activating IP_3_-induced calcium release (IICR).

IP_3_Rs are a family of calcium channels involved in a variety of cellular functions. There are three different IP_3_R isoforms encoded by separate genes. The three IP_3_Rs share a high degree of sequence homology and are found in a variety of tissues including the heart (6, 7). Cardiac IP_3_Rs are implicated in regulating the progression of cardiac hypertrophy (8, 9). Within the cardiomyocyte, IP_3_Rs are known to localize in the dyadic cleft, sarcoplasmic reticulum and at the outer/inner nuclear membrane (1, 8–10). Several lines of evidence have also implicated nuclear calcium transients as a significant contributor to cardiomyocyte hypertrophy. Nuclear or perinuclear IP_3_Rs may promote nuclear-restricted calcium release events that initiate gene transcription (10). Nuclear calcium transients are involved in the activation of transcription factors such as histone deacetylase 5 (HDAC5) (1, 11). However the mechanism by which cardiomyocytes can discriminate between calcium signals from ECC and calcium signals that target gene transcription it still unclear, as calcium release events mediated by ECC are also efficiently transmitted to the nuclear matrix. Hypertrophic agents such as ET-1 also increase contractility (9, 12), which may afford a mechanism for decoding IP_3_-dependent signals without requiring subcellular compartment-specific IP_3_R activation.

The IP_3_R-2 isoform is considered the predominant isoform in the heart (13, 14). Transgenic IP_3_R-2 rodent models have either supported (15, 16), or contradicted (17, 18) the role of IP_3_R channels in cardiac hypertrophy. As such, it is still unclear whether IP_3_R channels are significant contributors to cardiac physiology and pathologic remodeling such as hypertrophy (19). It has been shown that all three IP_3_R isoforms, at least at the mRNA level, are expressed in the heart of humans and mice (8). This opens the question of whether IP_3_R-1 and −3 are able to functionally compensate for IP_3_R-2 deficiencies in these models.

We now show that all three IP_3_R isoforms are expressed in cardiomyocytes and that they are essential for the progression of ventricular hypertrophy induced by ET-1. IP_3_R-dependent activation of the hypertrophic program was not dependent upon nuclear-specific calcium transients, but instead was mediated by increased contractility induced by ET-1. Lastly, these results were independent of increased IP_3_R expression both in vitro and in vivo.

## Results

### Expression of IP_3_R protein in neonatal and adult ventricular cardiomyocytes

Previous studies have indicated that IP_3_R-2 mRNA is predominant compared to other IP_3_R isoforms in ventricular cardiomyocytes (13, 14). Using highly specific antibodies we analyzed the protein expression and localization of the three IP_3_R isoforms in primary neonatal and adult rat cardiomyocytes. In neonatal ventricular cardiomyocytes all three IP_3_R isoforms were expressed throughout the cell (Fig.1 A-C, second column). Expression was more prominent in the perinuclear region consistent with previous reports (1). Similarly, all three IP_3_R isoforms are expressed throughout the cell in adult rat cardiomyocytes (Fig.1 D-F, second column). In contrast to neonatal ventricular cardiomyocytes, IP_3_R expression in adult cardiomyocytes was more equally distributed throughout the cell with no obvious concentration in the perinuclear region. This expression pattern overlaps with SERCA2 localization indicating is it present throughout the sarcoplasmic reticulum in both neonatal and adult cardiomyocytes (Fig. S1A, C). As expected, IP_3_R localization was distinct from the localization with RYR ((1); Fig. S1B, D). These results indicate that in addition to IP_3_R-2, IP_3_R isoforms type 1 and 3 are expressed in rat neonatal and adult ventricular cardiomyocytes.

**Fig 1.**
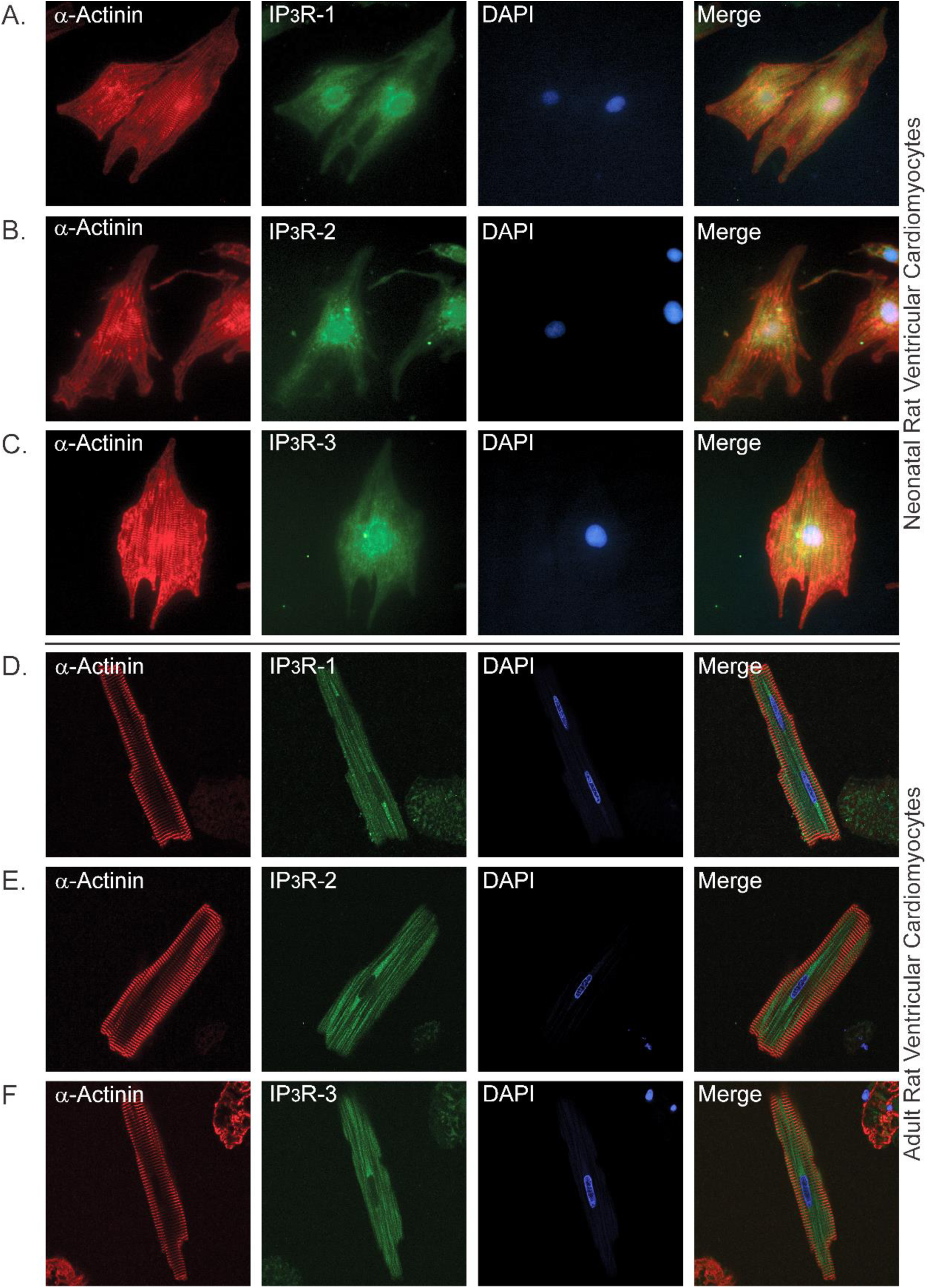
Expression and distribution IP_3_R isoforms in primary neonatal and adult ventricular cardiomyocytes. Immunofluorescence staining of neonatal (rows A-C) and adult (rows D-F) ventricular cardiomyocytes. Column 1 is stained with α-actinin to label sarcomeres. Column 2 is stained with indicated IP_3_R antibodies. Column 3 is DAPI staining of the nucleus. Column 4 is the merged images of each row.

### Altered contractility induced by ET-1 is attenuated by decreasing IP_3_R expression

ET-1 is known act as a positive inotropic agent in cardiomyocytes (9, 12). We monitored intracellular calcium in spontaneously contracting neonatal ventricular cardiomyocytes treated with ET-1. As shown in Fig. 2A (control bars) there is an increase in the calcium oscillation frequency in neonatal cardiomyocytes stimulated with 100 nM ET-1. Previous studies using overexpression of IP_3_ 5’-phosphatase to inhibit IP_3_ signaling suggested that IP_3_R channels do not play a role in this response to ET-1 (1). To evaluate the role of IP_3_R channels in the process more specifically, we used siRNA knockdown of each individual IP_3_R isoform. Western blot analysis confirmed efficient and specific knockdown of each of the three IP_3_R isoforms (Fig. S2). Knockdown of individual IP_3_R isoforms did not have significant effects on the increased oscillation frequency induced by ET-1 stimulation (Fig. 2A). Next we used double knockdowns (IP_3_R 1/2, 2/3 and 1/3) and determined their effect on the response to ET-1. Double knockdown of IP_3_R 1 and 2 resulted in a partial suppression of the response to ET-1. Triple knockdown of all three IP_3_R isoforms completely suppressed the response of cardiomyocytes to ET-1 (Fig. 2A). Thus, the increased contractility in response to ET-1 requires IP_3_R activity, and furthermore all three IP_3_Rs are expressed and play a functionally redundant role in the response to ET-1 stimulation.

**Fig 2.**
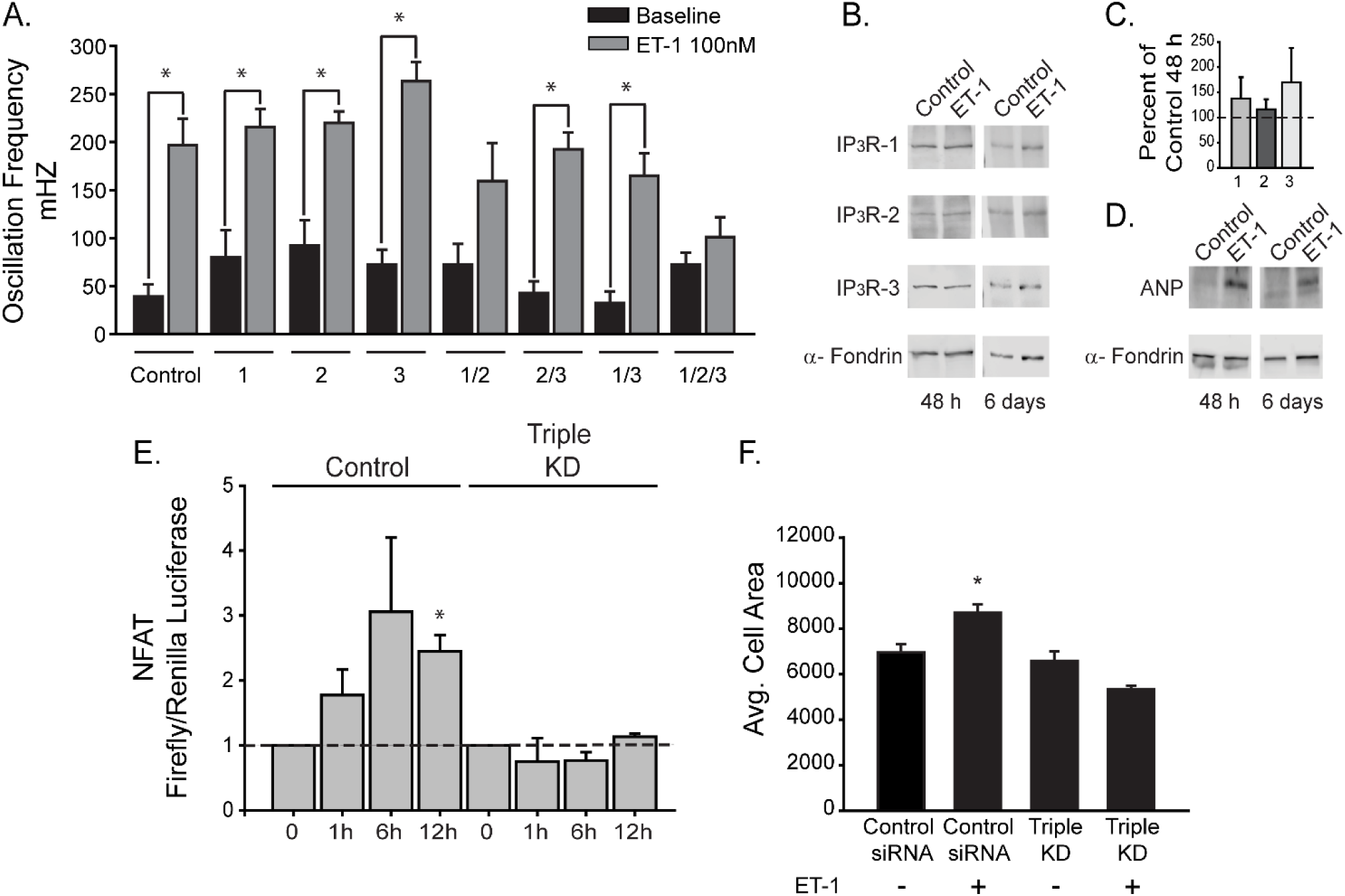
All three IP_3_R isoforms contribute to hypertrophic signaling by ET-1. (A) Neonatal cardiomyocytes were transfected with control or IP_3_R siRNA as indicated. Cells were treated with ET-1 and oscillation frequency was quantified from at least 4 separate experiments (control n=6, 1 n=6, 2 n=7, 3 n=6, 1/2 n=6, 2/3 n=5, 1/3 n=5 and 1/2/3 n=4). (B). Western blot of indicated proteins after 48 hours or 6 days treatment with 100 nM ET-1. (C). Quantification of IP_3_R levels after ET-1 treatment for 48 hours expressed as percent of control. (D) Atrial natriuretic peptide (ANP) levels after 48 hours and 6 days treatment with ET-1. Blotting with alpha-Fodrin was used as control. (E) NFAT activity using luciferase reporter construct after ET-1 treatment for the indicated times in control siRNA and triple IP_3_R siRNA transfected cardiomyocytes. (F) Average cell area before and after 48 hours treatment with ET-1. Neonatal cardiomyocytes were transfected with control siRNA and triple IP_3_R siRNA. All data are shown as mean ± SEM, *P < 0.01 vs control.

### Attenuating IP_3_R expression inhibits ET-1 induced NFAT activation and hypertrophy

At the cellular level cardiac hypertrophy is characterized by activation of the hypertrophic transcriptional program by NFAT ultimately leading to increased cardiomyocyte cell size. ET-1 is one of the best described neuro-hormonal factors known to induce cardiomyocyte hypertrophy (10, 16). As mentioned previously, IP_3_R mRNA levels are altered during hypertrophy (7, 13), however the effect on IP_3_R protein levels is not clear. In order to assess whether IP_3_R protein expression is altered during ET-1 induced hypertrophy, we treated neonatal cardiomyocytes with ET-1 for 48 hours and 6 days. Using this paradigm, protein expression of all three IP_3_R isoforms is unchanged compared to non-treated controls (Fig. 2B-C) despite robust induction of the hypertrophic marker protein atrial natriuretic peptide (ANP; Fig. 2D). It has also been shown that IP_3_R mRNA levels are altered in both animal models (9, 17) and in the human failing heart (8). We examined IP_3_R expression level in left ventricular heart tissue from one control and two end stage heart failure patients. Protein expression of all three IP_3_R isoforms could be readily detected in the three human heart samples. We found no significant differences in the protein expression level of IP_3_R isoforms in human failing hearts relative to control (Fig. S3 and Supplementary Table 1).

Next we determined whether IP_3_R expression is essential for activation of the hypertrophic program downstream of ET-1 stimulation. Treatment of cells with ET-1 led to significant NFAT activation within 12 hours as determined by a luciferase reporter assay (Fig. 2E). Triple knockdown of all three IP_3_R isoforms completely abrogated NFAT activation in response to ET-1 stimulation (Fig. 2E), a finding consistent with our observation that triple IP_3_R knockout suppresses the increased calcium oscillation frequency in response to ET-1 (Fig. 2A). Treatment of cardiomyocytes with ET-1 leads to an increase in cell size, the cardinal feature of hypertrophy ((1, 5); Fig. 2F). Triple knockdown of all three IP_3_R isoforms completely abrogates the increased cell size induced by to ET-1 treatment (Fig. 2F; S4). Thus, all three IP_3_Rs contribute to signaling downstream of ET-1 in cardiomyocytes and are essential for activation of the hypertrophic program.

### Calcium release induced by ET-1 is not restricted to the nuclear compartment

Rhythmic contraction of the heart is mediated by intracellular RyR calcium channels in a process termed calcium-induced calcium release (CICR). An important question is how IP_3_R-mediated calcium transients could be “decoded” in the constant background of CICR in the beating heart. One prevalent model is that nuclear-compartment specific IP_3_R calcium transients mediate gene expression in response to hypertrophic agents (10). However, as CICR calcium transients also diffuse into the nucleus the specific mechanism is not clear. Furthermore, NFAT is a cytosolic protein in the inactive state, and translocates to the nucleus in response to elevations in cytosolic calcium (5). To resolve these discrepancies, we used the genetically encoded calcium indicator GCaMP6s targeted to the nucleus in order to measure nuclear calcium transients unambiguously (Fig. 3A-B). Using the spectrally separated indicator Fura-2 to measure cytosolic calcium, we were able to quantify calcium release in both nuclear and cytosolic compartments with high specificity. As shown in Fig. 3C-D, calcium release was detectable in both compartments of spontaneously beating cardiomyocytes. After ET-1 stimulation there is an increase in the oscillation frequency consistent with the positive inotropic effects of ET-1 (Fig. 3C-D), which we showed is absolutely dependent upon IP_3_R expression (Fig. 2A). Visual observation of Fura-2 versus H2b-GCaMP6s traces revealed essentially overlapping plots (Fig. 3C-D), however the H2b-GCaMP6s signal had lower resolving power at higher oscillation frequencies. Expression of cytosolic GCaMP6s revealed that this was due to a buffering effect of the indicator, not an intrinsic property of the nuclear matrix (Fig. S5). In order to compare the cytosolic versus nuclear oscillations more quantitatively, we calculated the oscillation frequency over the course of the experiment in 20 second bins in both the cytosolic and nuclear compartments (Fig. 3E). Data points above the line in Fig. 3E would indicate a higher frequency in the nucleus (thus highlighting nuclear only events). Conversely, data points below the line are representative of events that occur in the cytosol but were not detectable in the nucleus. Presented in this way, we were unable to visualize any “nuclear only” release event either before or after ET-1 stimulation. We did, however, note a few “cytosol only” events at various frequencies. This may be due to either a cytosol restricted transient or insufficient sensitivity of the GCaMP6s sensor. Regardless, if there are nuclear restricted calcium transients they are either rare or low amplitude events. The results presented herein are also entirely consistent with the large body of evidence indicating that NFAT is activated by changes in calcium release frequency, not amplitude (20–22).

**Fig 3.**
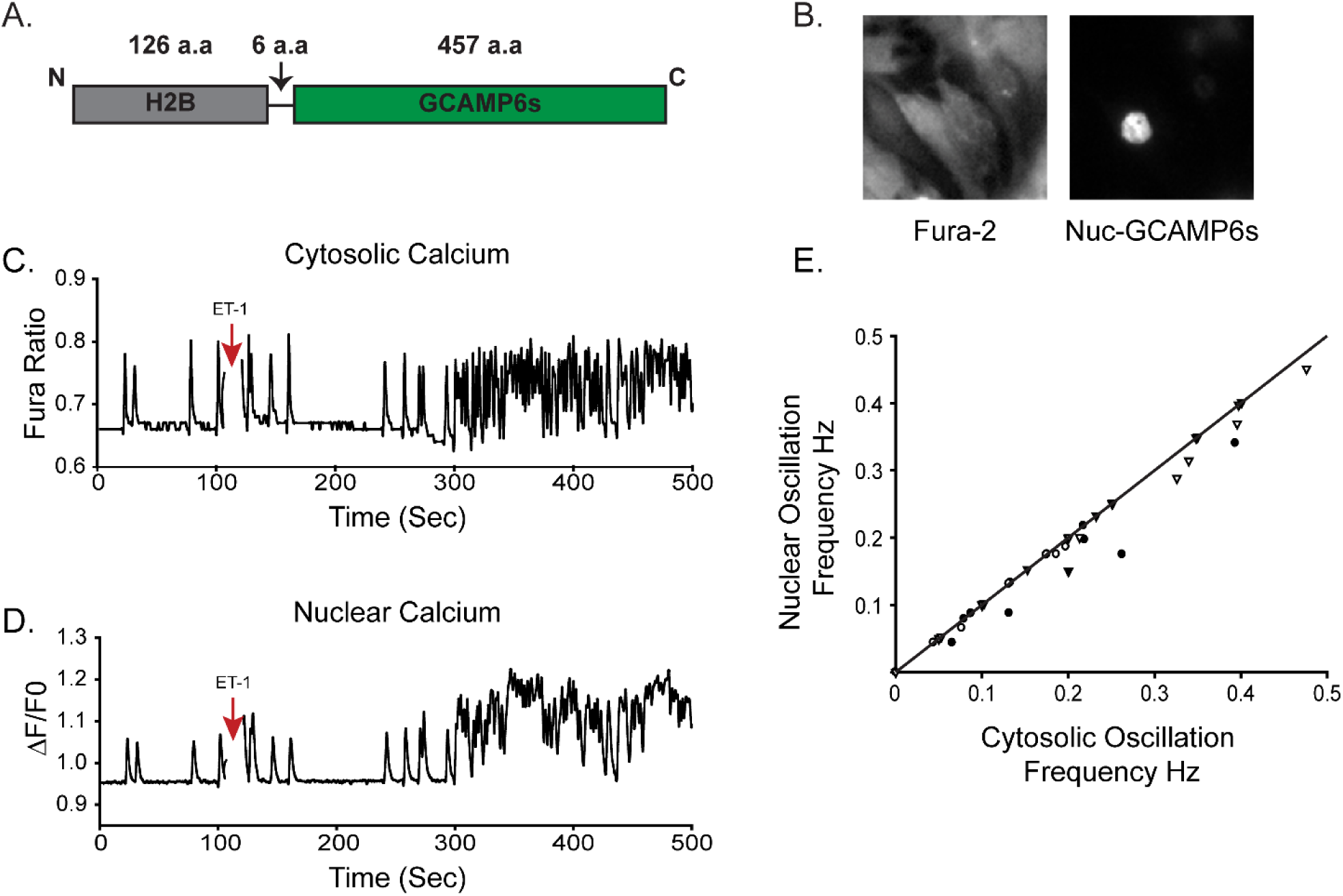
Nuclear and cytosolic calcium in response to ET-1 stimulation. (A) Schematic cartoon of H2BGCaMP6s. (B) Representative image of neonatal cardiomyocytes expressing H2B-GCaMP6s loaded with Fura-2 AM. (C) Single cell imaging of a neonatal cardiomyocyte treated with 100 nM ET-1 at indicated time. (D) H2b-GCaMP6s signal in the same cell as in C. (E) Plot of cytosolic vs. nuclear oscillation frequency. Each symbol represents a single coverslip averaging 5-10 cells for a total of four separate coverslips. Frequency data was quantified from 20 seconds bins.

## Discussion

IP_3_Rs likely play a key role in the progression of hypertrophy (1, 9, 16). However, the majority of the studies on the role of IP_3_Rs in hypertrophy have focused solely on IP_3_R-2, despite the fact that the other two isoforms are also expressed in the heart (8, 17). We show here that all three IP_3_R channels are expressed at readily detectable levels in cardiomyocytes in both rodent and human heart. Furthermore, all three channels contribute to calcium release and activation of the hypertrophic program in a functionally redundant manner. Lastly, our findings indicate that IP_3_Rs contribute to activation of the hypertrophic program after ET-1 stimulation primarily by mediating the increase in calcium release frequency/contractility. We do not find evidence, at least in this model, for nuclear-specific calcium events.

Our finding that IP_3_Rs play functionally redundant roles in ventricular cardiomyocytes may resolve some controversies regarding transgenic models of IP_3_R function in the heart. Previously, the IP_3_R-2 was thought to be the predominant isoform expressed in the heart. Consequently, most of the studies that focus on the role of IP_3_R in the heart have focused solely on IP_3_R-2 function (13, 23). Global genetic knockout of the IP_3_R-2 in mice does not cause any significant difference in the hypertrophic response in pressure overload or dilated cardiomyopathy mouse models (17). However, the potential involvement of IP_3_R-1 and IP_3_R-3 activity has not been investigated further. Another group has showed that global knockout of IP_3_R-2 eliminates the positive inotropic effects of endothelin-1 (ET-1) in the atria and protects against arrhythmias (15). Overexpression of the ligand binding domain of the IP_3_R inhibits signaling through the channel by buffering IP_3_ levels (so called “IP_3_ sponge” (24)). Targeted inducible overexpression of the IP_3_ sponge in the heart inhibited hypertrophy in response to isoproterenol and angiotensin-II (16). As the IP_3_ sponge would inhibit signaling through all three IP_3_R channels, these findings are consistent with our results suggesting that all three IP_3_Rs can contribute to hypertrophic signaling in cardiomyocytes.

The subcellular localization and precise aspects of how IP_3_Rs regulate spatio-temporal aspects of calcium release in the contracting myocyte is an area of debate. In particular, how IP_3_R signals are decoded to regulate processes such as transcriptional activation in the beating cardiomyocyte is unclear. IP_3_Rs are known to be primarily localized at the ER/SR membranes in most cells, however in the heart it is thought to be concentrated at the nuclear and perinuclear membranes, where it is thought to play a key role in gene transcription (1, 10, 11, 25). Nuclear localized IP_3_R would thus facilitate spatially restricted calcium transients to the nuclear matrix. It has been shown that IP_3_ and ET-1 can trigger nuclear calcium sparks and nuclear localized calcium transients (10, 25–27), however it is unclear whether these transients originate from the cytosol/perinuclear area. We show now that all three IP_3_Rs are expressed throughout the SR of the cardiomyocyte in both isolated rat cardiomyocytes and human tissue. Using nuclear-localized GCaMP6s to unambiguously monitor nuclear calcium, our results simultaneously imaging nuclear and cytosolic calcium indicate that nuclear-only calcium transients, if present, are an exceedingly rare events. Rather, our results indicate unequivocally that IP_3_Rs mediate the positive inotropic effects of ET-1 in a functionally redundant manner. The increase calcium release frequency would then result in activation of NFAT and activation of the hypertrophic program.

## Materials and Methods

### Antibodies, expression constructs, and reagents

Rabbit polyclonal antibody against type-1 IP_3_R was developed in-house and is specific for the type-1 isoform (28). The rabbit polyclonal antibody against type-2 IP_3_R have been described elsewhere (6) and was kindly provided by Dr. Richard Wojcikiewicz (SUNY Upstate). Mouse monoclonal anti-IP_3_R type-3 was purchased from BD Bioscience. Mouse monoclonal anti-α-actinin and anti-ryanodine receptor antibody was purchased from Sigma-Aldrich. Mouse anti-SERCA2 antibody was from ThermoFisher and rabbit anti-ANP was purchased from Abcam. Anti-alpha-fodrin was purchased from EMD Millipore. Secondary antibodies conjugated to Alexa-488 and Alexa-555 were from Molecular Probes, and peroxidase-conjugated antibodies were from Jackson ImmunoResearch. Expression constructs 9X NFAT-apha-MHC-Luc was a gift from Jeffery Molkentin (Addgene plasmid # 51941), pGP-CMV-GCaMP6s was a gift from Douglas Kim ((29); Addgene plasmid # 40753) and Tol2-elavl3-H2B-GCaMP6s was a gift from Misha Ahrens (Addgene plasmid # 59530). Endothelin-1 was purchased from Bachem. Silencer pre-design siRNAs were purchased from Ambion. Fura-2 AM was purchased from Molecular Probes, and the dual luciferase reporter assay kit was from Promega. All other reagents were purchased from Sigma-Aldrich.

### Preparation of primary neonatal cardiomyocytes

Neonatal rat ventricular cardiomyocytes (NRVM) were obtained from 1 to 2 day old Sprague-Dawley rat hearts as previously described, with minor modifications (30). Cardiomyocytes were plated into fibronectin-coated culture dishes and incubated at 37°C in 5% CO2 incubator. Two days after plating, media was replaced with 50% Ham's F10 −50% DMEM culture medium with β-Darabinofuranoside (ARA-C; 1 μM) to inhibit growth of fibroblasts. NRVMs were transfected using Lipofectamine 3000 following manufacturer's instructions. All experiments were carried out 48 hours after transfection. All vertebrate animal procedures were approved by the Animal Welfare Committee (AWC) at UTHealth.

### Preparation of adult cardiomyocytes

Calcium-tolerant adult rat ventricular myocytes (ARVM) were isolated from hearts of wildtype Sprague-Dawley rats (300 – 320 g) as described by Louch et. al. (31). Briefly, animals were anesthetized with chloralhydrate (400mg/kg b.w. i.p.) and heparinized (5,000 U/kg b.w.) via direct injection into the vena cava inferior. The hearts were aseptically removed and directly placed in ice-cold Krebs-Henseleit (KH) buffer (133.5 mM NaCl, 4 mM KCl, 1.2 mM NaH2PO4, 10 mM HEPES, 1.2 mM MgSO4, 10 mM BDM) containing glucose (5.5 mM) before being perfused with a Langendorff preparation. Perfusion (3 min) with KH buffer at 37°C lacking Ca2+ was followed by perfusion with recirculating KH buffer containing 2% BSA (wt/vol), 50 μM Ca2+ and type II collagenase. After 20 minutes of perfusion, hearts were minced, and undigested tissue was separated with a 230 μm mesh sieve. The cell suspension was allowed to settle with gravity within 5 to 7 min, and the cell pellet was re-suspended in KH containing 2% BSA (wt/vol), and calcium was slowly reintroduced to a final concentration of 1 mM. Cardiomyocytes were plated and culture under 5% carbon dioxide.

### Immunofluorescence labeling of isolated cardiomyocytes

Adult and neonatal cardiomyocytes were plated at a density of 300 cells per mm^2^ on glass coverslips coated with fibronectin. On day 4 the medium was exchanged and the cells were treated with 100 nM ET-1 for 48 hrs. Cells were fixed with 4% paraformaldehyde in PBS. Briefly, cells were incubated with either rabbit polyclonal anti-IP_3_R-1 (1:250), rabbit polyclonal anti-IP_3_R-2 (1:250), mouse monoclonal anti-IP_3_R-3 (1:250), mouse monoclonal anti-α-actinin (1:250), mouse anti-SERCA2 (1:250), or mouse anti-ryanodine2 (1:250), overnight at 4°C. Followed by incubation with secondary antibodies conjugated to Alexa-488 and Alexa-555 for 1 hr.

### Immunofluorescent labeling of human heart failure samples

Disease heart tissue was obtained from patients undergoing heart transplantation due to advanced heart failure. Immediately after explant the tissue is flash frozen with liquid nitrogen for future analyses. Control heart tissue was obtained from organs that were declined for transplantation due to non-cardiac reasons. Frozen left ventricular cardiac tissues were cryo-sectioned onto charged glass slides. The sections were fixed with 4% paraformaldehyde in PBS. Tissue was stained with anti-IP_3_R-1 (1:100), anti-IP_3_R-2 (1:100), anti-IP_3_R-3 (1:100), or anti-α-actinin (1:100), for 1hr at 37°C. Subsequently, slides were washed and incubated with secondary antibodies conjugated to Alexa-488 and Alexa-555 for 1 hr. All experiments on human samples were approved by the Institutional Review Board (IRB) of UTHealth.

### Cell size determination

NRVM were plated on glass coverslips. Two days after plating cells were transfected with control siRNA or triple IP_3_R siRNA. Following transfection cells were treated with ET-1 for 48 hours. After treatment cells were fixed with 4% paraformaldehyde in PBS. Subsequently, coverslips were stained with anti-α-actinin and secondary antibodies conjugated to Alexa-555. For measurement of cell area, at least 30 fields randomly chosen were analyzed in each coverslip. Cardiomyocytes area was measured in captured images using ImageJ software.

### Western Blotting

Cells were harvested by gently scraping plates with a cell scraper and washing once with cold PBS. Cell lysis buffer (150 mM NaCl, 50 mM Tris-HCl at pH 7.8, 1% Triton X-100 and 1 mM EDTA) was added to the cell pellet. Samples were cleared of insoluble debris by centrifugation at 20,000g at 4ᵒC. Cell lysates were quenched with SDS sample buffer. Samples were resolved by SDS–PAGE and analyzed by Western blotting. Where indicated, cardiomyocytes were treated 100 nM ET-1 for 48 hours or six days.

### Calcium imaging

NRVM were plated on fibronectin-coated glass coverslips and were transfected with triple IP_3_R siRNA targeting rat IP_3_R-1, IP_3_R-2 and IP_3_R-3. The total amount of siRNA transfected was 12.5 pmol per 350,000 cells. Transfected cells were identified by co-transfection with cDNA for YFP (0.25pmol per 350,000 cells). Cells were imaged after 48 hrs. For imaging, cardiomyocytes were incubated with 5 μM Fura-2 AM in imaging solution (1% BSA, 107mM NaCl, 20 mM HEPES, 2.5 mM MgCl2, 7.5 mM KCl, 11.5mM glucose, and 1 mM CaCl2) for 30 min at RT. The solution was replaced with imaging solution without Fura-2 AM for an additional 20 min. Images were acquired using a Nikon TiS inverted microscope as previously described (32). Responses to 100 nM ET-1 were recorded on YFP only positive cells. In order to specifically look at nuclear calcium transients we used GCaMP6s fused to a sequence encoding human histone H2b at the 5′ end (33). We sub-cloned H2b-GCAMP6s into the mammalian expression vector pcDNA 3.1(+).NRVM were transfected with different siRNAs (IP_3_R-1, IP_3_R-2 or IP_3_R-3) and with H2b-GCAMP6s two days after plating. Cells were loaded with Fura-2 AM in imaging solution 48 hrs after transfection. Responses to 100nM ET-1 were acquired at 1 Hz during continuous recording. Oscillation frequency was determined manually. An oscillation was counted when the Fura-2 ratio rose 10% above the baseline ratio. Similar to Fura-2, an oscillation was counted when GCaMP6s fluorescence rose above 5% from baseline fluorescence.

### NFAT luciferase

For assessment of NFAT activation cells were co-transfected with 9xNFAT-TATA luciferase plasmid (34) and pRL-TK control vector (2:1). All experiments were performed 48 hours after transfection. Cells were harvested and cell extracts were assayed using dual luciferase reporter assay as specified by manufacturer's protocol (Promega). All data are shown as mean ± SEM, with statistical significance determined at p < 0.01 vs control using an unpaired Student's t-test.

## Acknowledgements

We would like to thank Dr. Richard Wojcikiewicz (SUNY Upstate) for his generous gift of IP_3_R-2 antisera. We would also like to thank Dr. Shane Cunha (McGovern Medical School at UTHealth) for advice regarding primary cardiomyocyte preparation. Lastly, we would like to thank Ann-Bin Shyu (McGovern Medical School at UTHealth) for assistance with the luciferase assays. This work was support by NIH grants R01GM081685 (DB) a Research Supplement to Promote Diversity in Health-Related Research on the same grant (to MIG), R01HL61483 (HT), Friede Springer Herz Stiftung (AK), and the Roderick MacDonald Research Fund 15RDM005 (AK). This work was also supported by startup funds provided by the McGovern Medical School at UTHealth (DB).

## Supplemental Figure Legends

**Fig S1.**
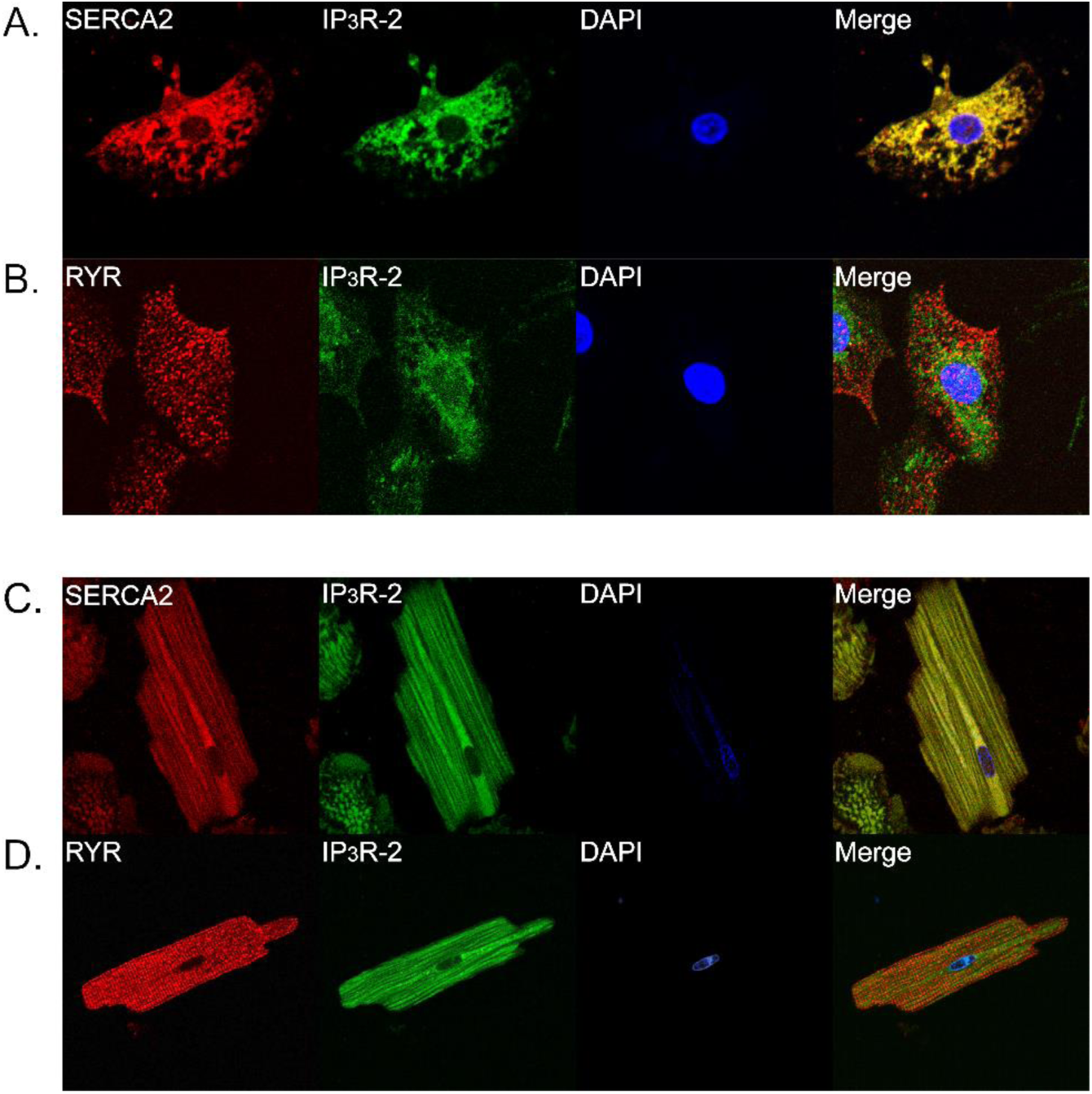
Expression and distribution of IP_3_R, SERCA2, and RYR in primary ventricular cardiomyocytes. Immunofluorescence staining of neonatal ventricular cardiomyocytes stained with SERCA2 and IP_3_R-2 (A); RYR2 and IP_3_R-2 (B). Immunofluorescence staining of adult ventricular cardiomyocytes with SERCA2 and IP_3_R-2 (C); RYR2 and IP_3_R-2 (D).

**Fig. S2.**
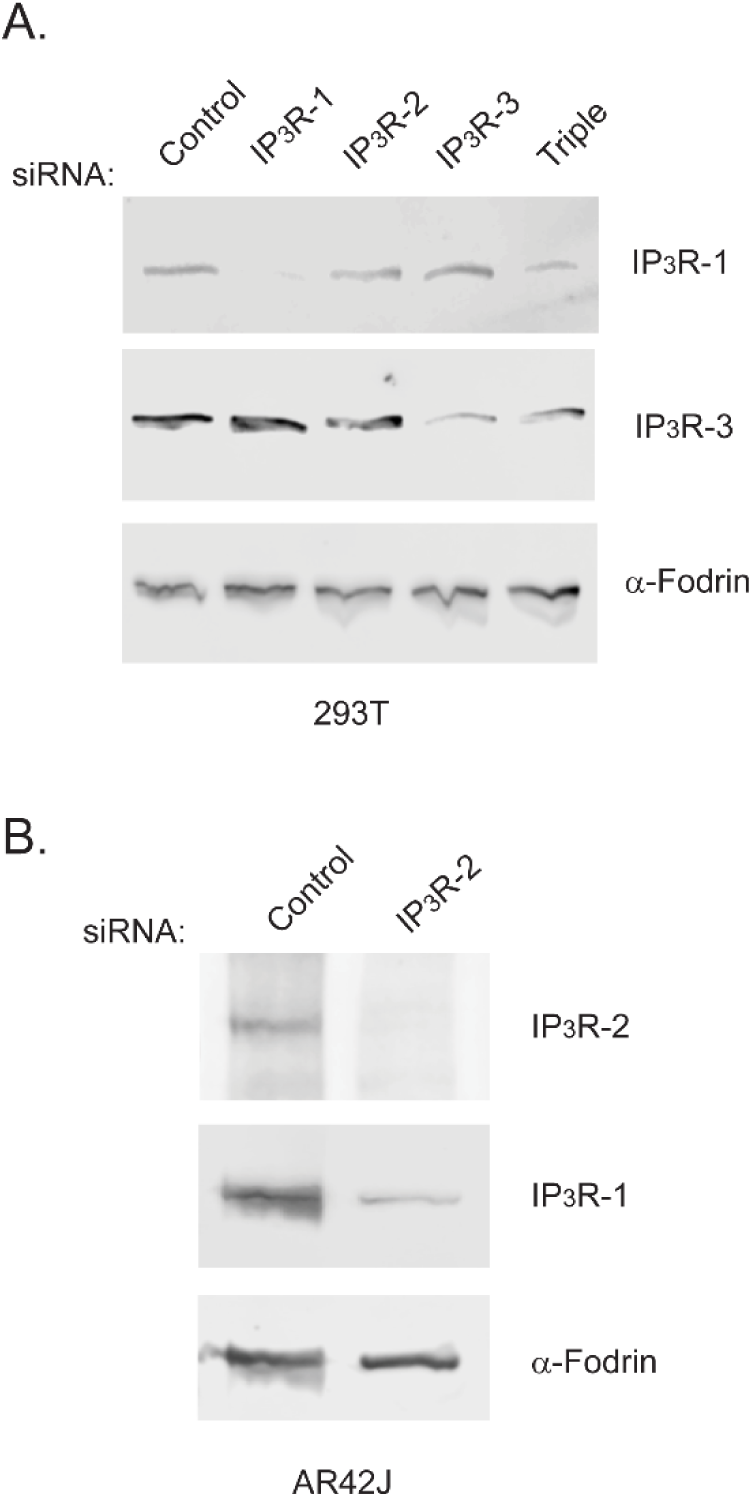
siRNA-mediated knockdown of IP_3_R-1, −2 and −3. Western blot analysis of 293T (A) and AR42J (B) cell lines transfected with control siRNA and siRNA oligos specific for each IP_3_R isoform as indicated. IP_3_R-1 siRNA significantly reduced endogenous IP_3_R-1 expression in 293T where control siRNA did not. Similarly IP_3_R-3 siRNA significantly reduced IP_3_R-3 expression. Both siRNAs were specific for the individual isoforms. Triple IP_3_R siRNA transfection inhibited the expression of both IP_3_R-1 and −3 in 293T cells (A). IP_3_R-2 expression was below detection levels in 293T cells. To probe the efficiency of IP_3_R-2 siRNA, we used AR42J cells which express high amounts of this isoform (B). IP_3_R-2 siRNA completely inhibited IP_3_R-2 expression, and also partially reduced IP_3_R-1 levels. IP_3_R-3 expression was below detection levels in AR42J cells. In both panels, alpha-fodrin was used as loading control.

**Fig. S3.**
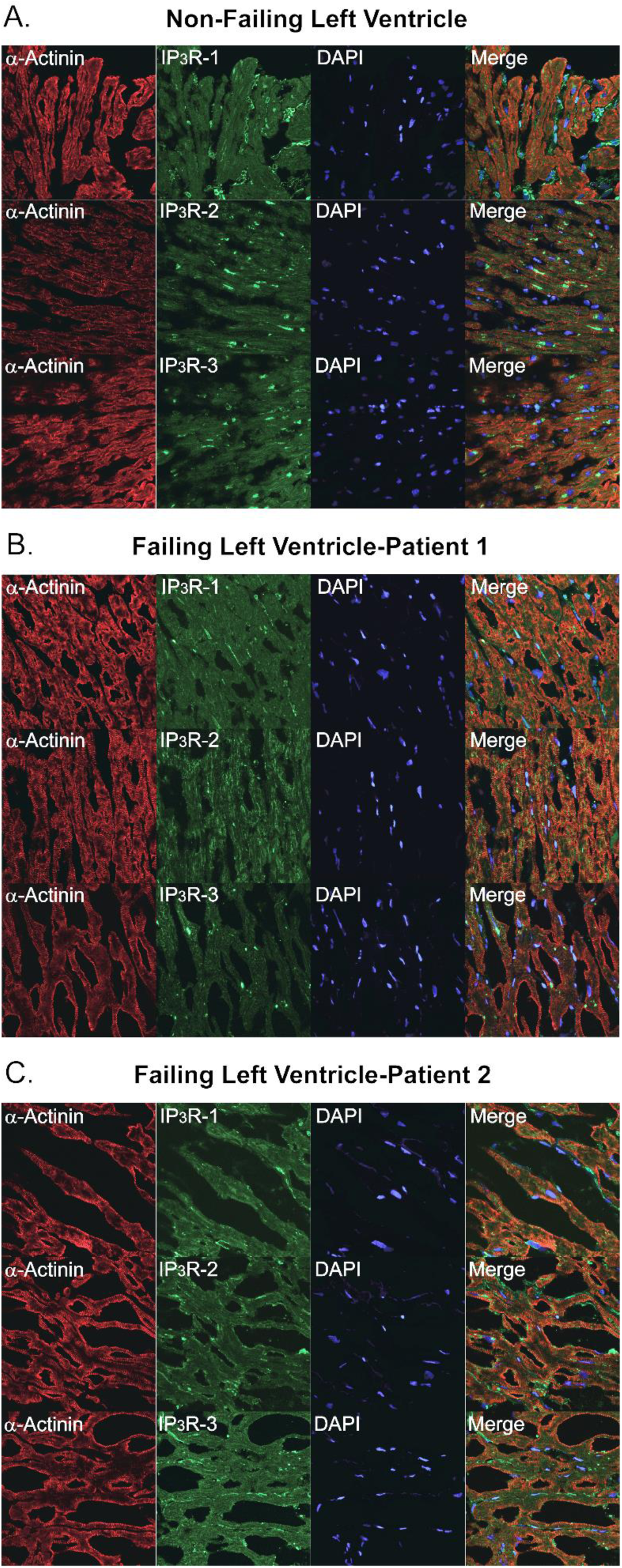
Expression of IP_3_R isoforms in non-failing and end stage heart failure samples. Immunofluorescence staining of non-failing human left ventricle (A). Left ventricular heart failure patient 1(B) and patient 2 (C). Column 1 is stained with α-actinin to label sarcomeres. Column 2 is stained with indicated IP_3_R antibodies. Column 3 is DAPI staining of the nucleus. Column 4 is the merged images of each row.

**Fig. S4.**
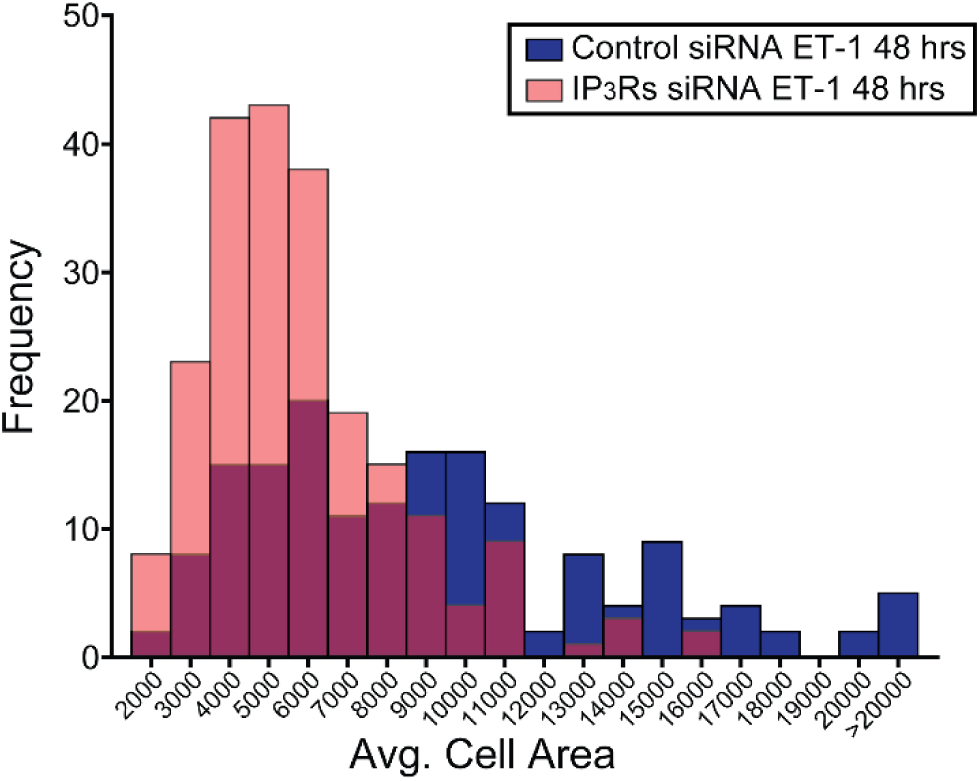
Histogram of cardiomyocyte size distribution after ET-1 stimulation. Histogram of neonatal ventricular cardiomyocytes cell size distribution showing control (n=166) and triple IP_3_R siRNA (n=218) transfected cells treated with ET-1 for 48 hrs.

**Fig S5.**
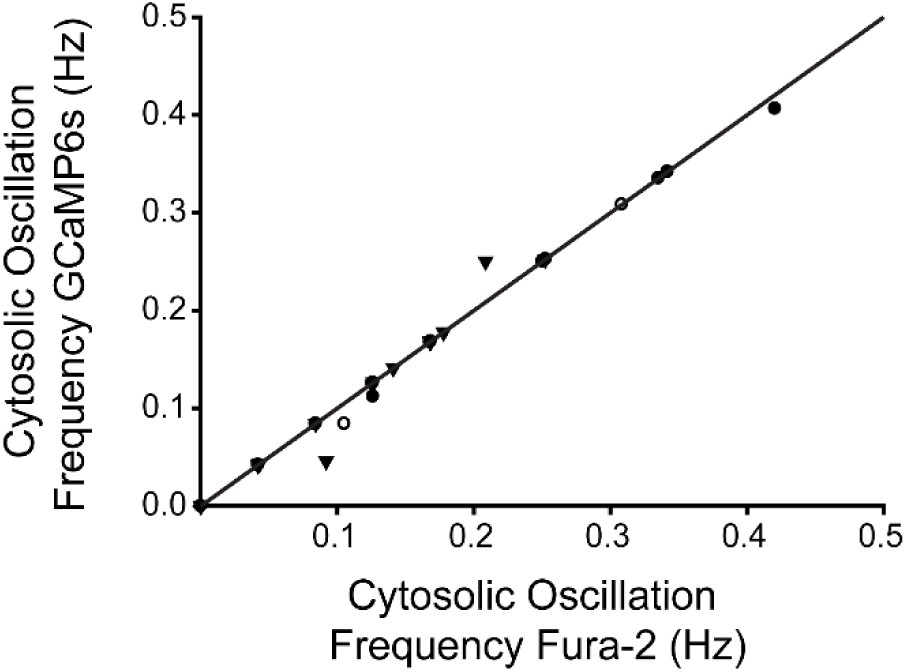
Plot of cytosolic GCaMP6s versus Fura-2 oscillation frequency. Cytosolic GCaMP6s was imaged simultaneously with cytosolic Fura-2 oscillation in neonatal cardiomyocytes and relative oscillation frequencies from each indicator were quantified. Each symbol represents a single coverslip averaging 5-10 cells for a total of three separate experiments. Frequency data was quantified from 20 seconds bins.

**Supplementary Table 1.** Clinical characteristics of patient samples. Control sample was obtained from a heart that was declined for transplantation due to non-cardiac reasons. Patient 1 and 2 samples were obtained from patients that suffered with end stage heart failure. NYHA indicates New York Heart Association; Stage 1 hypertension, systolic pressure ranging from 140 to 159 mm Hg or a diastolic pressure ranging from 90 to 99 mm Hg; LVEDD, left ventricular end diastolic diameter-normal range 4.2-5.9 cm; EF, Ejection Fraction, normal EF ranges from 55-70%; LVPWd, Left ventricular posterior wall end diastole and end systole-normal range 0.6-1.1 cm.

|  | Control | Patient 1<br>HF | Patient 2<br>HF |
| --- | --- | --- | --- |
| NYHA Class | 0 | 4 | 3.5 |
| Age | 43 | 57 | 64 |
| Sex | F | M | M |
| Hypertension | - | Stage 1 | Stage 1 |
| LVEDD | - | 7.2 | 6.7 |
| LVPWd | - | 1.4 | 1.3 |
| EF | 60% | 20% | 20% |

